## Supplementary Information for "Clinical application of tumour-in-normal contamination assessment from whole genome sequencing"

Jonathan Mitchell, Salvatore Milite, Jack Bartram, Susan Walker, Nadezda Volkova, Olena Yavorska, Magdalena Zarowiecki, Jane Chalker, Rebecca Thomas, Luca Vago, Alona Sosinsky, Giulio Caravagna

Corresponding: (AS); (GC).

#### **Supplementary Figures**

Supplementary Figure S1. *In silico* validation of TINC.
Supplementary Figure S2. CNA profiles of lung cancer samples.
Supplementary Figure S3. Sequential gating/sorting strategies for monitoring MRD.
Supplementary Figure S4. Application of TINC to ALL patients.
Supplementary Figure S5. Normal sample source proportions for the haematological cohort.
Supplementary Figure S6. PASS and FAIL status for the haematological cohort.
Supplementary Figure S7. Normal sample source and TINC status for the sarcoma cohort.
Supplementary Figure S8. Hotspot somatic variants being at risk of subtraction.
Supplementary Figure S9. Correlation between TIN score and VAF for CHIP mutations.
Supplementary Figure S10-S13. TINC analysis reports.

#### **Supplementary Tables**

Supplementary Table 1. Requirements for the collection of samples from haematological malignancies

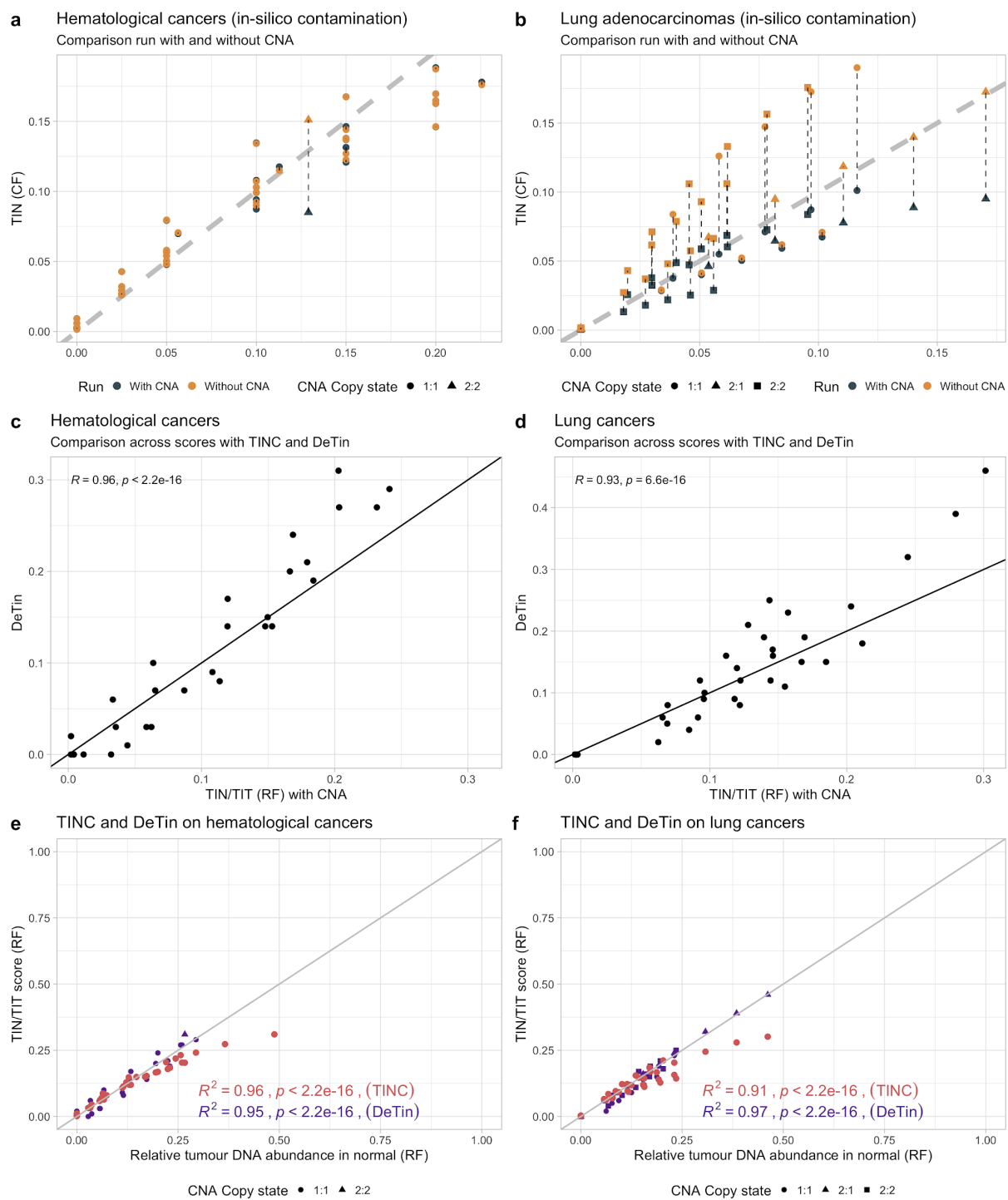

**Supplementary Figure S1. *In silico* validation of TINC.** **a,b** *In silico* tests with TINC run with and without CNA data, for the two cohorts presented in the Main Text. The x-axis reports contamination in units of cell fractions (CF). The solutions are connected by vertical lines. **c,d** Correlation between DeTin and TINC scores (stat\_cor R function; Pearson method with two-sided p-value and squared correlation coefficient), computed as ratios to follow DeTin convention. All the statistics were computed using the Pearson method with two-sided p-value and squared correlation coefficient. **e,f** Extended version of Figure 2d,e, to

include cases with very high contamination levels - beyond the parameters that are acceptable for the flowchart in Figure 4. All the statistics were computed using the Pearson method with two-sided p-value and squared correlation coefficient. Source data are provided as a Source Data file.

**A**

**Comparative CNA**

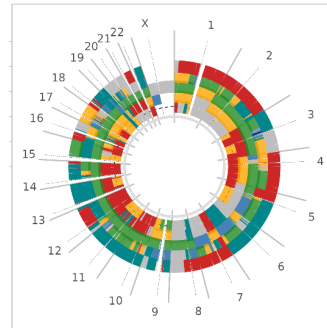

**B**

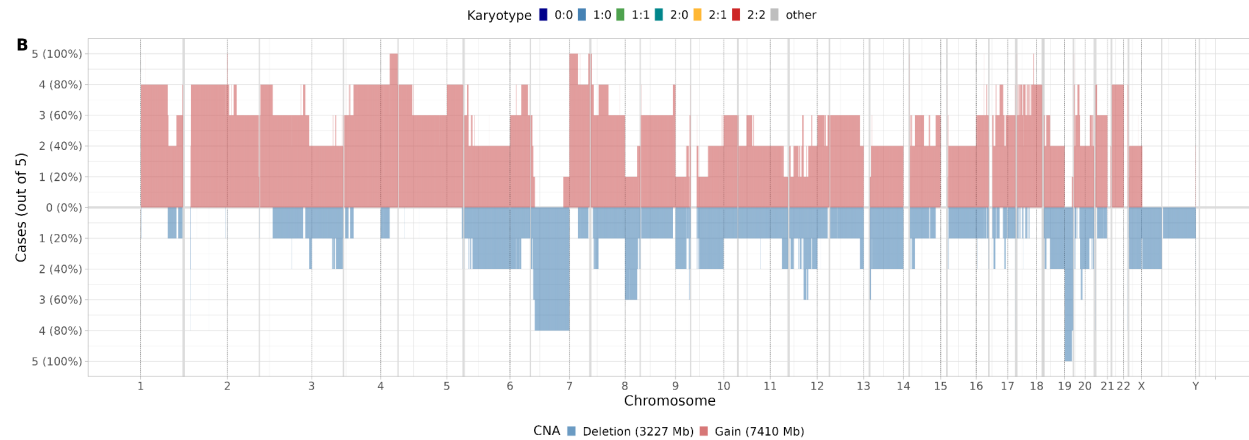

**Supplementary Figure S2. CNA profiles for the lung cancer samples used in this study.** **a** Circos plot of allele-specific CNA data for the lung cancer samples used in Figure 2c. Colours correspond to different allele-specific segment values across the whole genome. **b** Distribution of deletions and gains in cases from panel a. On average, 78% of the tumour genome in each of the samples is affected by clonal copy number events.

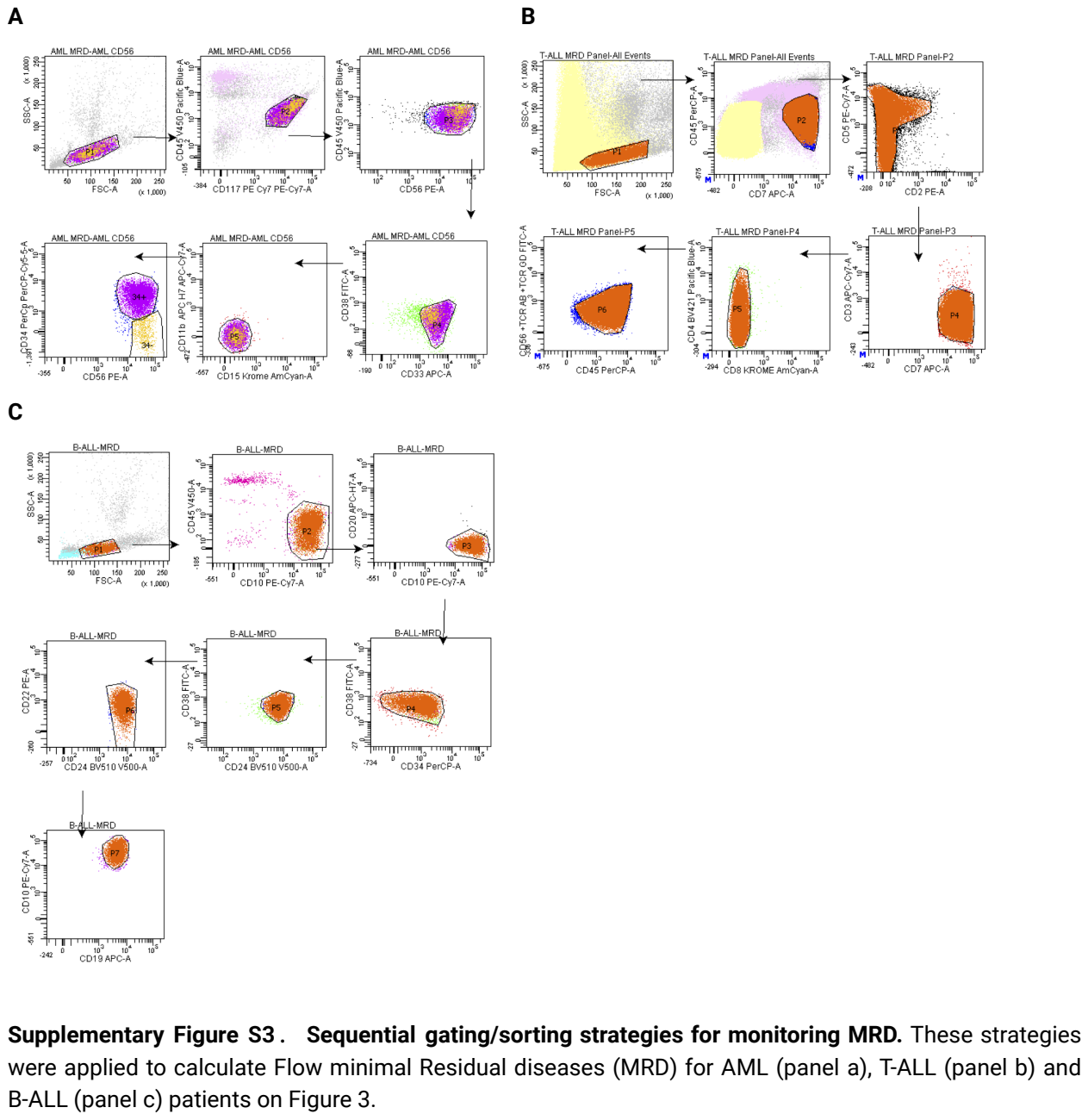

Supplementary Figure S3. Sequential gating/sorting strategies for monitoring MRD. These strategies were applied to calculate Flow minimal Residual diseases (MRD) for AML (panel a), T-ALL (panel b) and B-ALL (panel c) patients on Figure 3.

n = 67 ALL WGS (poor responders outside of the 100K Project)

Genomics England pipeline: n = 67 TINC with SNVs and CNA.

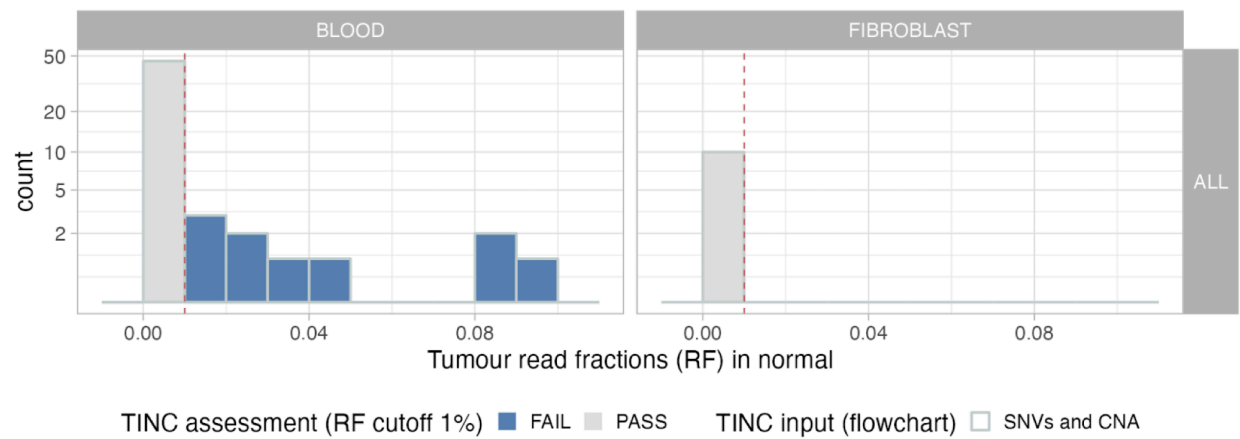

**Supplementary Figure S4. Application of TINC to ALL patients.** Contamination in tumour read fractions determined in $n = 67$ ALL patients whose normal samples do not fulfil recruitment standards for the 100,000 Genome Project, as reported in Supplementary Table 1. The samples are shown as in Figure 6. Note that blood normal samples from these cohorts show contamination in 11 out 46 cases. All 10 cases with fibroblasts as a source of normal samples show no sign of contamination. Source data are provided as a Source Data file.

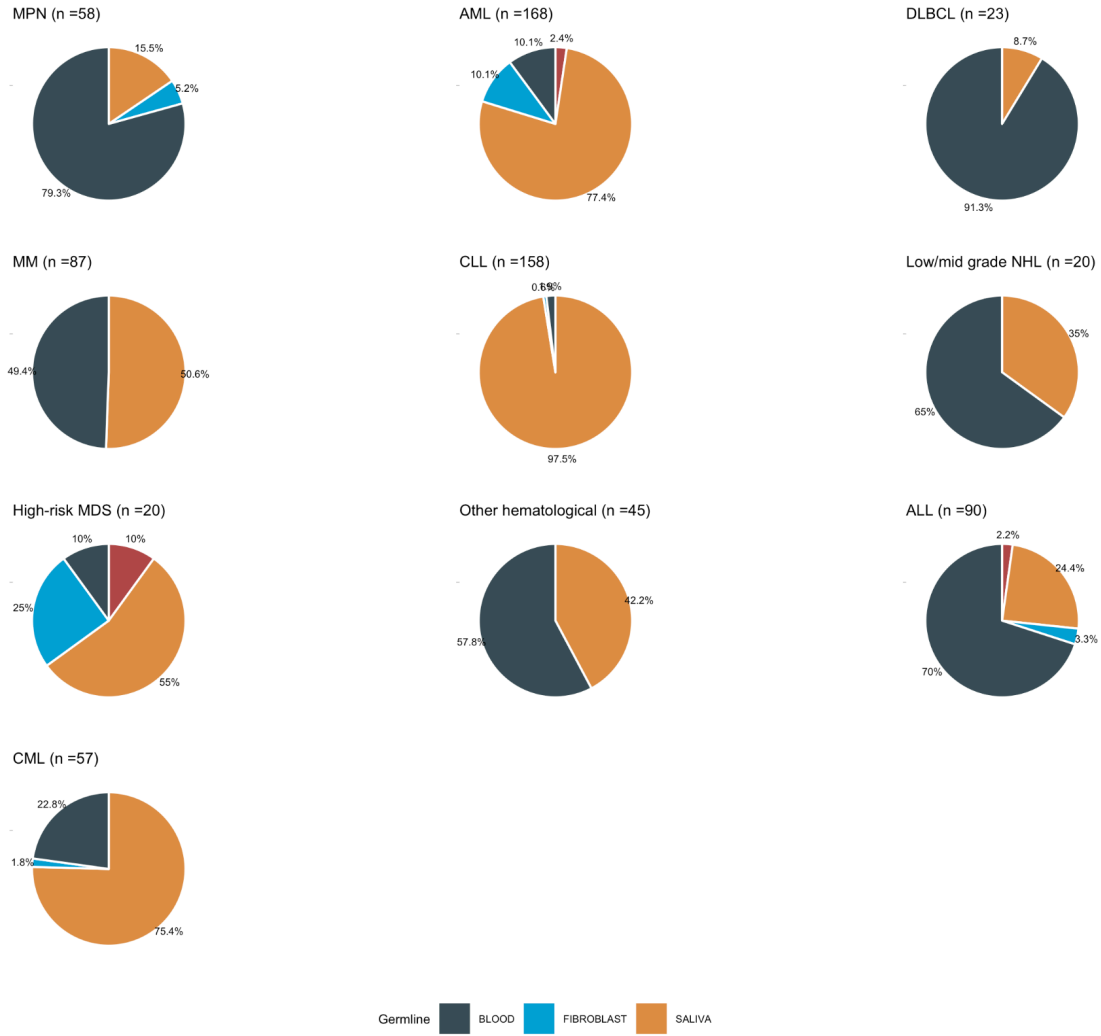

**Supplementary Figure S5. Normal sample source proportions for the haematological cohort.** Proportions for the haematological cohort shown in Figure 6 of the Main Text. Source data are provided as a Source Data file.

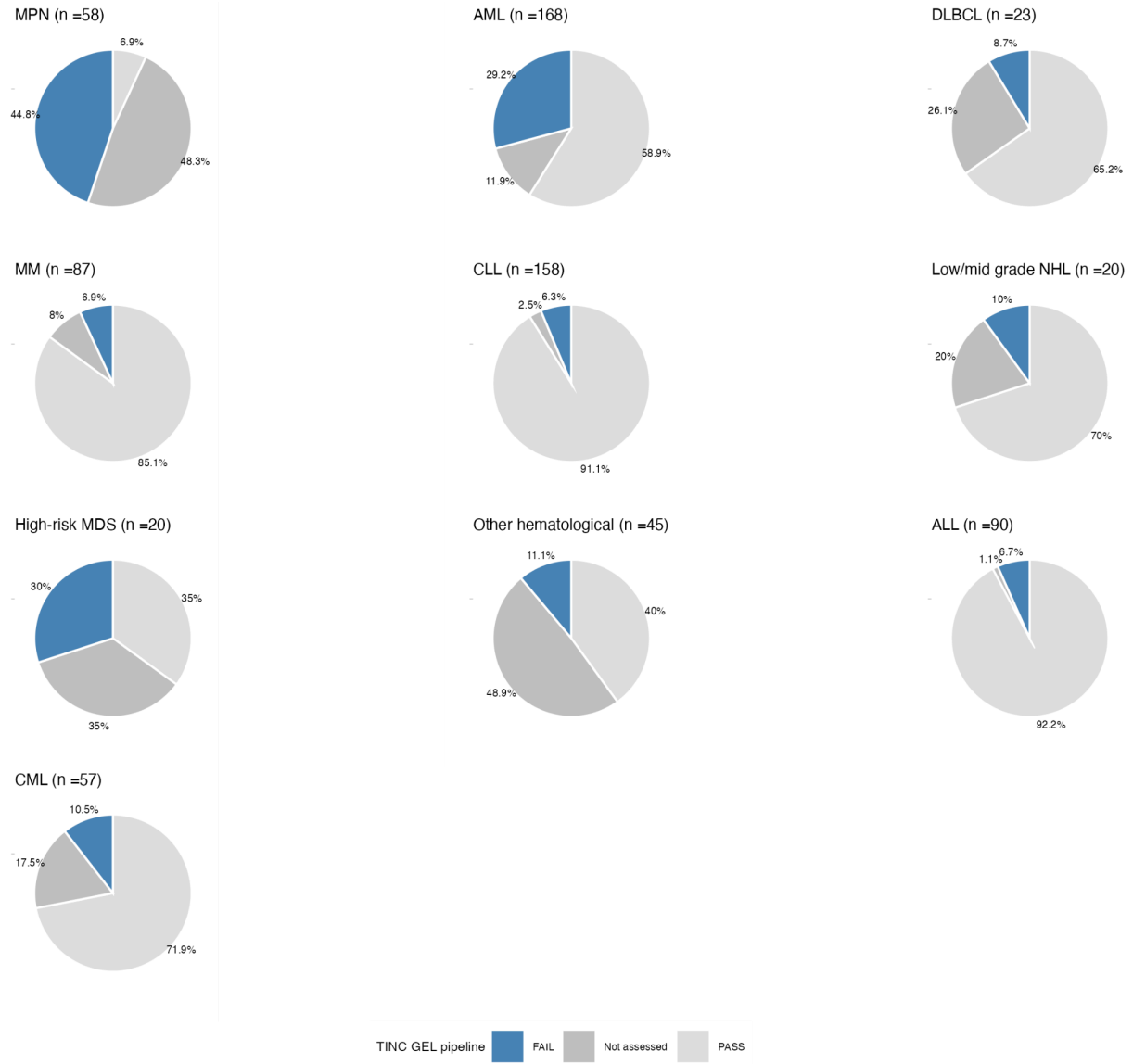

**Supplementary Figure S6. PASS and FAIL status for the haematological cohort.** The proportion of cases for the cohort shown in Figure 6 of the Main Text, that could not be analysed by Genomics England pipeline (tumour purity estimated to be below 25%) is shown in dark grey. Source data are provided as a Source Data file.

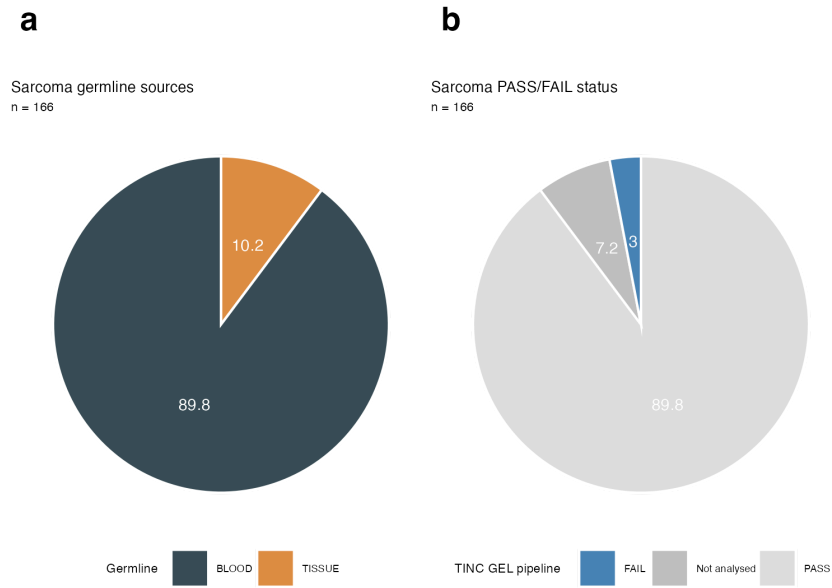

**Supplementary Figure S7. Normal sample source and TINC status for the sarcoma cohort. a** Normal sample source proportions for the cohort shown in Figure 6 of the Main Text. **b** Results from the Genomics England pipeline for the sarcoma cohort. Criteria used to PASS or FAIL samples are as in Figure 6. The proportion of cases that could not be analysed by Genomics England pipeline (tumour purity estimated to be below 25%) is shown in dark grey. Source data are provided as a Source Data file.

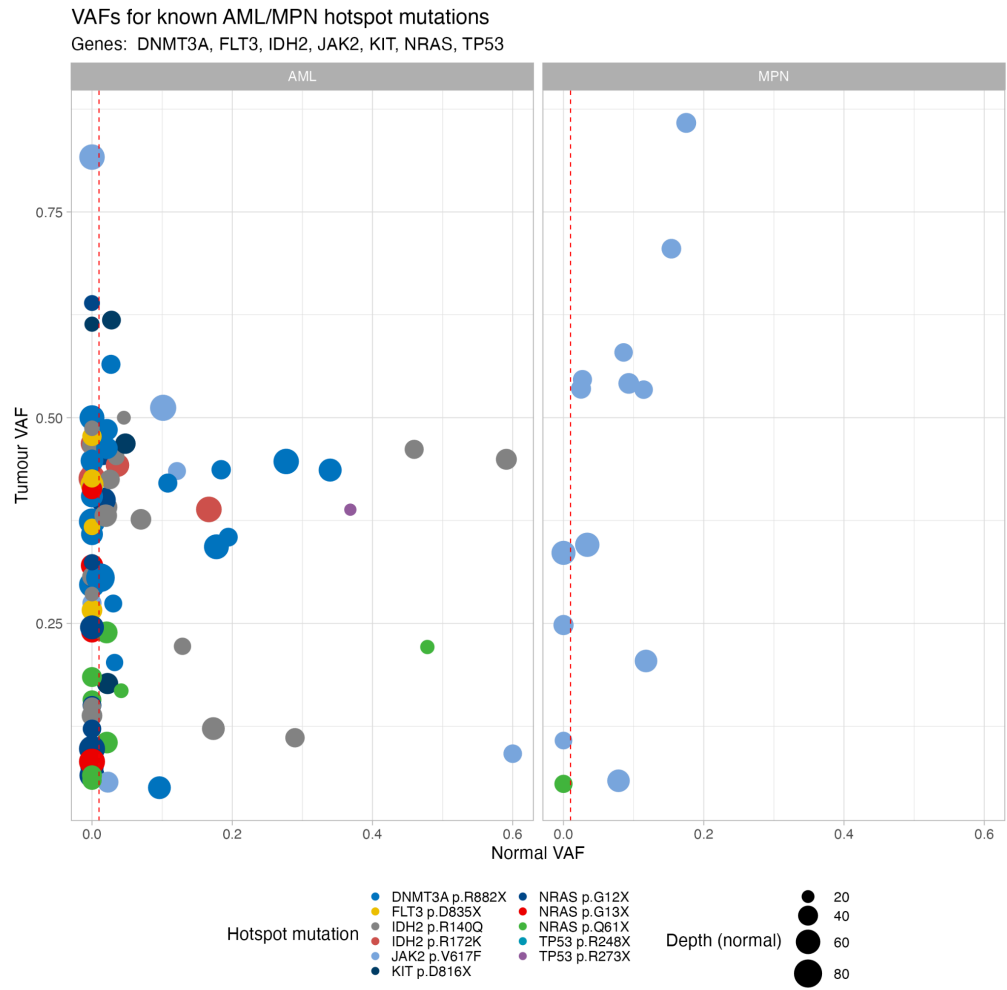

**Supplementary Figure S8. Hotspot somatic variants being at risk of subtraction.** Normal and tumour VAFs for a set of hotspot mutations in AML and MPN cohorts: *JAK2* (p.V617F), *FLT3* (p.D835X), *DNMT3A* (p.R882X), *TP53* (p.R248X, p.R273X), *KIT* (p.D816X), *NRAS* (p.G12X, p.G13X, p.Q61X) and *IDH2* (p.R140Q, p.R172K). The vertical dashed line denotes a 1% VAF in the normal.

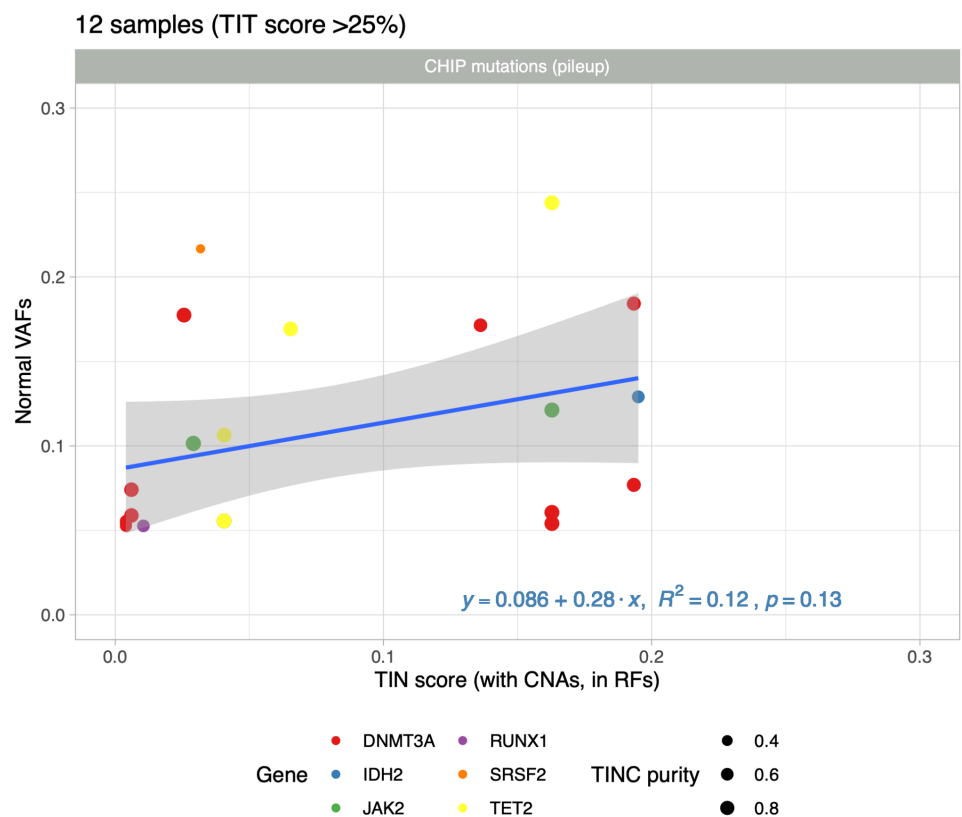

**Supplementary Figure S9. Correlation between TIN score and VAF for CHIP mutations.** Scatterplot of VAFs for CHIP-associated variants in the normal samples of 12 AML patients against TIN score (RF with SNVs and CNAs used in calculation). Size of the dot corresponds to the TIT score. Only samples with TIT>25% and variants with at least two supporting reads are shown. Smoothing performed using linear regression (stat\_cor R function; Pearson method with two-sided p-value and squared correlation coefficient). The shadowed area represents the 95% confidence level interval for predictions from a linear model.

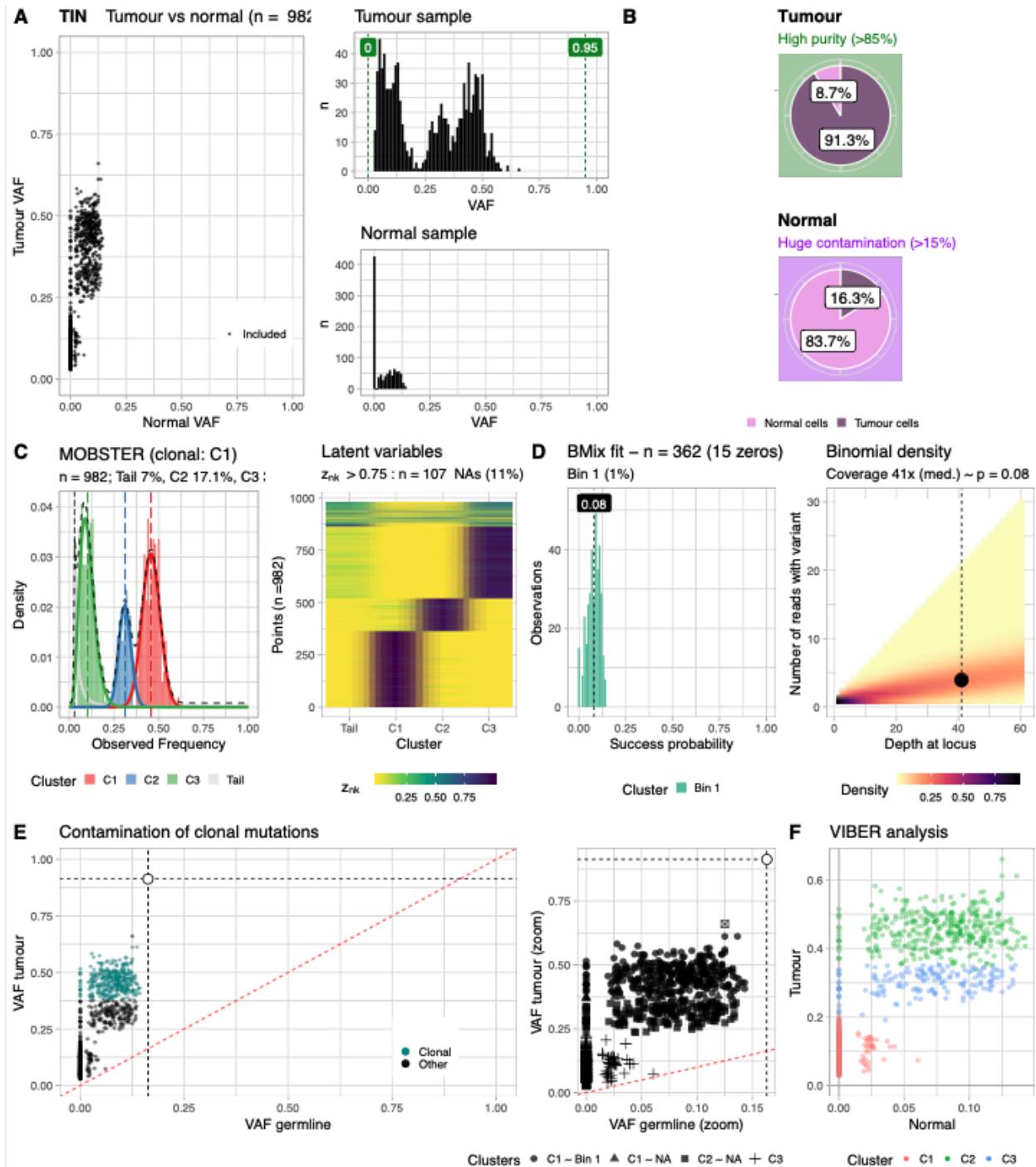

**Supplementary Figure S10. TINC analysis report (sample in Figure 7a-f).** a,b Data distribution for diploid SNVs, and TIT/TIN scores. c MOBSTER deconvolution and latent variables reporting the probability of each SNVs to be assigned to each one of the detected clusters. d BMix deconvolution and Binomial density (unitless) at the observed coverage. e Final assignments of clonal status to the input SNVs, with zoom for low-frequency variants. f Complementary VIBER multivariate clustering (cluster labelling is independent from MOBSTER clustering, panel c).

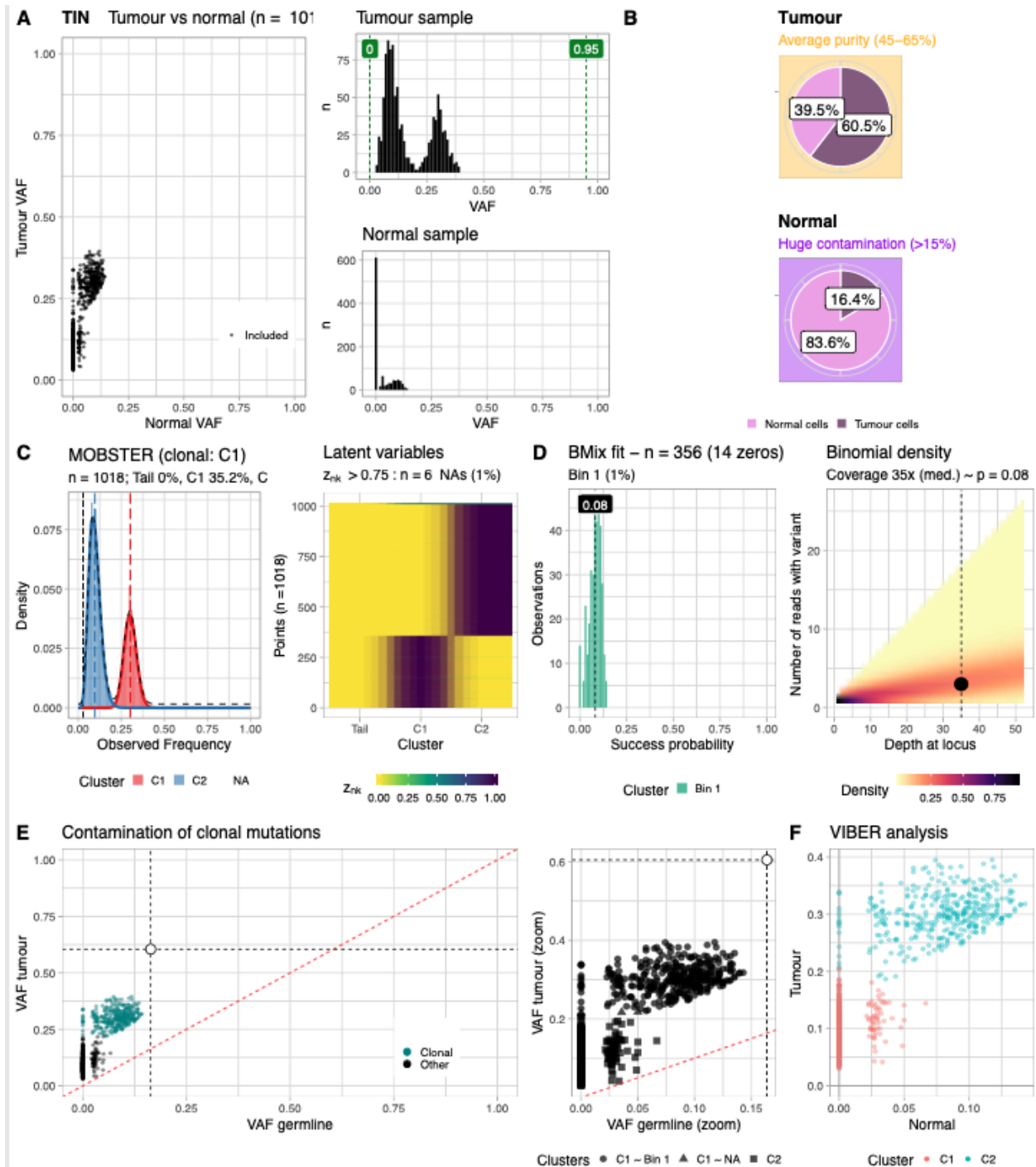

**Supplementary Figure S11. TINC analysis report (sample in Figure 7g-i).** **a,b** Data distribution for diploid SNVs, and TIT/TIN scores. **c** MOBSTER deconvolution and latent variables reporting the probability of each SNVs to be assigned to each one of the detected clusters. **d** BMix deconvolution and Binomial probability density (unitless) at the observed coverage. **e** Final assignments of clonal status to the input SNVs, with zoom for low-frequency variants. **f** Complementary VIBER multivariate clustering (cluster labelling is independent from MOBSTER clustering, panel c).

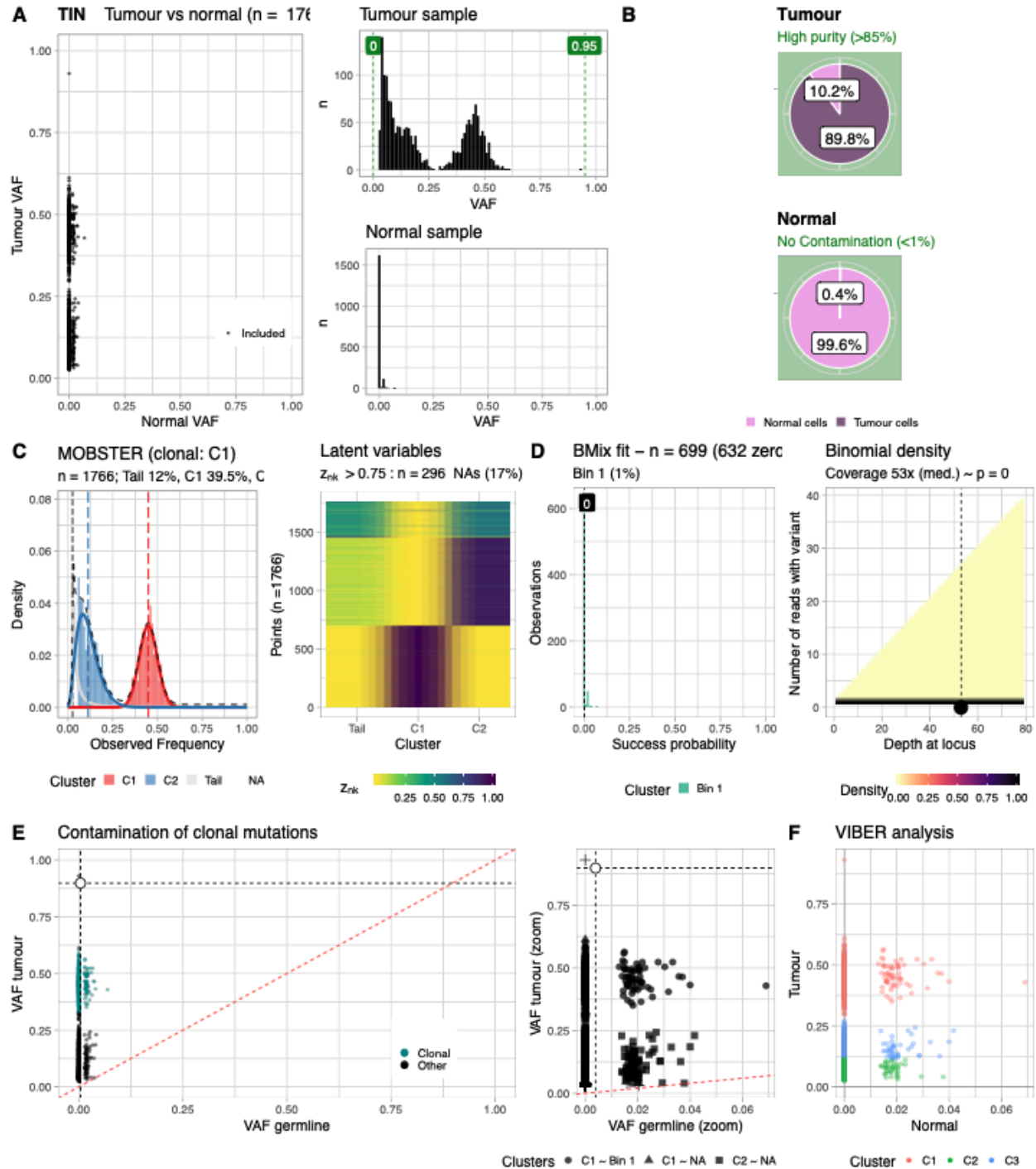

**Supplementary Figure S12. TINC analysis report (GEL sample without contamination).** a,b Data distribution for diploid SNVs, and TIT/TIN scores. c MOBSTER deconvolution and latent variables reporting the probability of each SNVs to be assigned to each one of the detected clusters. d BMix deconvolution and Binomial density (unitless) at the observed coverage. e Final assignments of clonal status to the input SNVs, with zoom for low-frequency variants. f Complementary VIBER multivariate clustering (cluster labelling is independent from MOBSTER clustering, panel c).

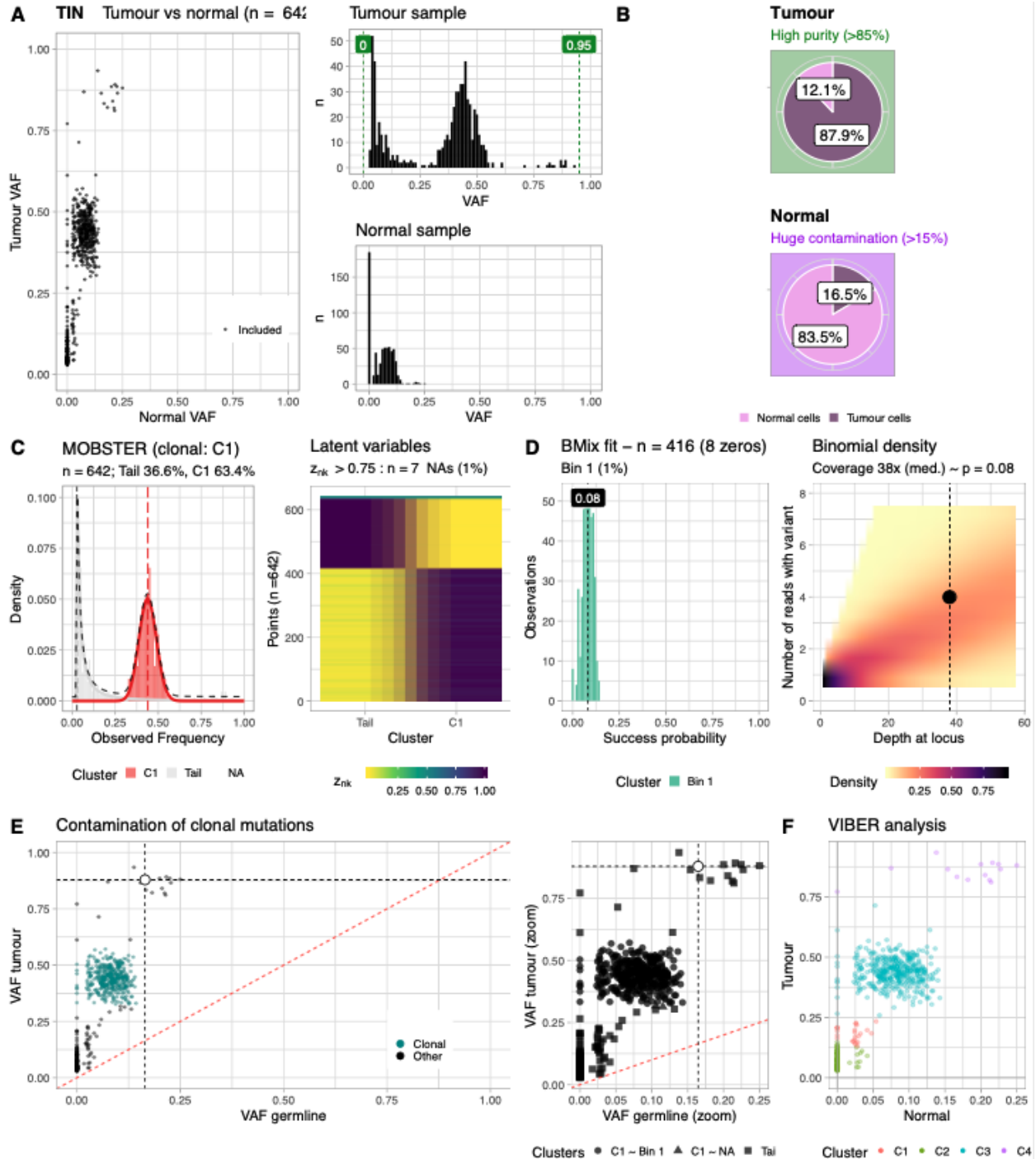

**Supplementary Figure S13. TINC analysis report (GEL sample with contamination, as in Figure 7). a,b** Data distribution for diploid SNVs, and TIN/TIT scores. **c** MOBSTER deconvolution and latent variables reporting the probability of each SNVs to be assigned to each one of the detected clusters. **d** BMix deconvolution and Binomial density (unitless) at the observed coverage. **e** Final assignments of clonal status to the input SNVs, with zoom for low-frequency variants. **f** Complementary VIBER multivariate clustering (cluster labelling is independent from MOBSTER clustering, panel c).

### Supplementary Tables

**Supplementary Table 1.**

Requirements for the collection of samples from haematological malignancies

| Clinical Indication | Normal sample source | Tumour sample source |
| --- | --- | --- |
| Acute Myeloid Leukaemia | Saliva<sup>a</sup><br>Cultured fibroblasts<br>Others<sup>b</sup> | Bone marrow aspirate or peripheral blood containing $\geq 20\%$ blasts morphologically or any blast percentage if there is an AML-defining genetic abnormality as per WHO 2016 Guidelines |
| Myelodysplastic Syndrome | Saliva<sup>a</sup><br>Cultured fibroblasts<br>Others<sup>b</sup> | Bone marrow aspirate or peripheral blood containing $\geq 5\%$ blasts morphologically |
| Chronic Myeloid Leukaemia<br>• Extreme 'Good' Responders<sup>c</sup> | Saliva<br>Peripheral Blood<sup>d</sup> | Bone marrow aspirate or peripheral blood (no minimum blast cell percentage required) |
| Chronic Myeloid Leukaemia<br>• Extreme 'Poor' Responders<sup>e</sup><br>• Additional Cytogenetic Abnormality<sup>f</sup><br>• Accelerated or Blast Phase<sup>g</sup> | Cultured fibroblasts<br>Others<sup>b</sup> | Bone marrow aspirate or peripheral blood (no minimum blast cell percentage required) |
| Unclassified Haematological Malignancies<sup>h</sup> | Saliva<sup>a</sup><br>Cultured fibroblasts<br>Others<sup>b</sup> | Bone marrow aspirate or peripheral blood (no minimum blast cell percentage required) |
| Acute Lymphoblastic Leukaemia | Saliva<sup>a</sup><br>Cultured fibroblasts<br>MRD negative peripheral<br>blood / bone marrow<br>aspirate<sup>i</sup> | Bone marrow aspirate or<br>peripheral blood containing<br>≥40% blasts<br>morphologically |
| Lymphoproliferative Disorders<sup>j</sup> | Saliva<br>Peripheral blood<sup>k</sup> | Fresh frozen tissue (i.e.<br>biopsy or resection)<br>Bone marrow aspirate or<br>peripheral blood containing<br>≥40% malignant cell nuclei<sup>l</sup><br>Other liquid sample<br>containing ≥40% malignant<br>cell nuclei<sup>m</sup> |
| Multiple Myeloma | Saliva<br>Peripheral blood<sup>n</sup> | CD138+ sorted cells with a<br>purity of ≥40%<sup>o</sup> |
| Chronic Lymphocytic Leukaemia | Saliva<sup>q</sup> | Bone marrow aspirate or<br>peripheral blood<sup>r</sup> containing<br>≥40% malignant cell nuclei |

###### Table notes

<sup>a</sup> Saliva is acceptable as a normal sample in myeloid malignancies only if sufficient treatment has been given to remove all circulating myeloid cells from the peripheral blood e.g. on day 5 after administration of two doses of anthracycline chemotherapy (or equivalent) in patients receiving intensive induction in Acute Myeloid Leukaemia.

<sup>b</sup> Alternative normal options are being pursued in the disease types indicated to facilitate recruitment to the programme including sorted CD3+ cells (T cells) and uncultured skin biopsies. If and when these normal sample types are acceptable, supplementary guidance will be issued detailing specific requirements.

<sup>c</sup> Extreme 'Good' Responders in Chronic Myeloid Leukaemia are defined as those patients who, after 3 months of treatment with a tyrosine kinase inhibitor, have achieved a *BCR-ABL* transcript level (by RQ-PCR) of <1% using International Standards.

<sup>d</sup> Peripheral blood is an acceptable source of normal DNA for patients who are classified as

Chronic Myeloid Leukaemia Extreme 'Good' Responders providing the *BCR-ABL* transcript level (by RQ-PCR) using International Standards is <0.1%.

<sup>e</sup> Extreme 'Poor' Responders in Chronic Myeloid Leukaemia are defined as those patients who, after 3 months of treatment with a tyrosine kinase inhibitor have a *BCR-ABL* transcript level (by RQ-PCR) of >10% using International Standards.

<sup>f</sup> Refers to any additional cytogenetic abnormality detected using karyotyping in the diagnostic sample in Chronic Myeloid Leukaemia other than a variant *BCR-ABL* transcript.

<sup>g</sup> Patients either presenting in Accelerated or Blast Phase Chronic Myeloid Leukaemia or progressing to Accelerated or Blast Phase Chronic Myeloid Leukaemia are eligible for recruitment.

<sup>h</sup> The definition of this category is broad but includes disorders such as 'Triple negative' Myeloproliferative Neoplasms (defined as no variant detected in *JAK2* exon 12, exon 14 (codon 617), *CALR* exon 9 or *MPL* exon 10), and Myelodysplastic/Myeloproliferative Overlap Syndromes.

<sup>i</sup> In Acute Lymphoblastic Leukaemia peripheral blood or bone marrow aspirate samples which are either negative for or have a diagnostic MRD marker (e.g. BCR or TCR gene rearrangement or *BCR-ABL* transcript) detectable at a level of <0.1% are suitable for use as the source of normal DNA.

<sup>j</sup> Any patient with a Lymphoproliferative Disorder (high or low grade) for which treatment is planned is eligible for recruitment to the project.

<sup>k</sup> Peripheral blood is suitable for use as the source of normal DNA in Lymphoproliferative Disorders providing there are no circulating tumour cells in the peripheral blood. Please note it is not necessary to undertake anything beyond normal standard of care assessments to demonstrate the absence of circulating tumour cells.

<sup>l</sup> Peripheral blood or bone marrow aspirate samples could be used as a source of tumour DNA in Lymphoproliferative Disorders providing the malignant lymphoid cells constitute $\geq 40\%$ of the nucleated cells in the sample.

<sup>m</sup> It is appreciated that there may be situations where malignant lymphoid cells constitute $\geq 40\%$ of the nucleated cells in a sample of a different body fluid e.g. pleural fluid; in these rare situations these would be an acceptable source of tumour DNA.

<sup>n</sup> Peripheral blood is an acceptable source of normal DNA in Myeloma providing there are no circulating plasma cells in the peripheral blood.

<sup>o</sup> All myeloma samples should undergo enrichment for CD138+ cells even if the starting plasma cell percentage of the bone marrow aspirate smear is $\geq 40\%$ in order to obtain the highest possible purity of plasma cells. Laboratories carrying out CD138+ cell enrichment / sorting will need to supply verification of the sorting technique and the CD138+ sorting checklist (Part 5: Appendix D) prior to commencement.

<sup>p</sup> It is appreciated that most myeloma samples will not have sufficient cells for tumour lysate collection for subsequent RNA extraction.

<sup>q</sup> Saliva collection in Chronic Lymphocytic Leukaemia should be postponed until such a time as the peripheral blood lymphocyte count is $<25 \times 10^9/L$.

<sup>r</sup> The lymphocyte count of peripheral blood samples to be used as the source of the tumour DNA in Chronic Lymphocytic Leukaemia should be $>25 \times 10^9/L$.
